# Bayesian inference for spatio-temporal stochastic transmission of plant disease in the presence of roguing: a case study to estimate the dispersal distance of Flavescence dorée

**DOI:** 10.1101/2022.12.14.520426

**Authors:** Hola Kwame Adrakey, Gavin J. Gibson, Sandrine Eveillard, Sylvie Malembic-Maher, Frederic Fabre

## Abstract

Estimating the distance at which pathogens disperse from one season to the next is crucial for designing efficient control strategies for invasive plant pathogens and a major milestone in the reduction of pesticide use in agriculture. However, we still lack such estimates for many diseases, especially for insect-vectored pathogens, such as Flavescence dorée (FD). FD is a quarantine disease threatening European vineyards. Its management is based on mandatory insecticide treatments and the removal of infected plants identified during annual surveys. This paper introduces a general statistical framework to model the epidemiological dynamics of FD in a mechanistic manner that can take into account missing hosts in surveyed fields (resulting from infected plant removals). We parameterized the model using Markov chain Monte Carlo (MCMC) and data augmentation from surveillance data gathered in Bordeaux vineyards. The data mainly consist of two snapshot maps of the infectious status of all the plants in three adjacent fields during two consecutive years. We demonstrate that heavy-tailed dispersal kernels best fit the spread of FD and that on average, 50% (resp. 80%) of new infection occurs within 10.5 (resp. 22.2) meters from the source plant. These values are in agreement with estimates of the flying capacity of *Scaphoideus titanus*, the leafhopper vector of FD, reported in the literature using mark–capture techniques. Simulations of simple control scenarios using the fitted model suggest that cryptic infection hampered FD management. Future efforts should explore whether strategies relying on reactive host removal can improve FD management.

**Author summary:** The dispersal of pathogen propagules is an important feature of spatial epidemiology that has a major impact on the incidence and distribution of disease in a population. In agriculture, properly characterising the dispersal of emerging disease is of great importance in designing science-based control strategies that allow pesticide use to be reduced. Although field epidemiological surveys can provide informative data, they are by nature rare while resulting from the interactions between disease spread and the undergoing surveillance and control. Here, we take advantage of a general statistical framework to model the epidemiological dynamics of Flavescence dorée (FD), a quarantine disease threatening European vineyards, in a mechanistic manner that can take into account missing hosts in surveyed fields (resulting from infected plant removals). We parameterized the model with a Bayesian approach using mainly two snapshot maps of the infectious status of all plants in three adjacent fields during two consecutive years. We demonstrate that on average, 50% (resp. 80%) of new FD infection occurs within 10.5 (resp. 22.2) meters of the source plant. Although FD mainly spreads locally from one year to the next, our results also indicate frequent long-distance dispersal events, a feature crucial to consider when designing control strategies.

## Introduction

The management of emerging plant pathogens is of foremost importance in the effort to reduce pesticide use in agriculture, particularly since the rate of introduction of alien species is higher than ever due to increasing globalisation [1]. The design of efficient management strategies has much to gain from the recent advances in epidemiological modelling. In particular, spatially explicit stochastic models are able to simulate realistic patterns of epidemic spread and test the effectiveness of a range of management strategies (for a review, see [2] and, more recently, [3–8]). However, for this approach to be feasible, the underlying model parameters must be estimated.

The main issue encountered when fitting spatially explicit mechanistic models is the type and nature of the available data. First, epidemiological surveys most often provide information on epidemic dynamics in the presence of control, which must accordingly be explicitly accounted for in models and fitting procedures [9,10]. Second, there is a trade-off between the number of fields that can be studied and the number of seasons over which they can be observed. The data collected most often contain only partial spatial and/or temporal information on the location of infected individuals. Moreover, the timing of true infection events is typically not observed. The first symptom expression in an individual plant (the most easily accessible variable during field surveys) can take place a long time after that plant becomes infected and even infectious. The latter situation results from the presence of cryptic infection, during which an infectious host is asymptomatic [2,11]. All of this context involves the existence of missing data. Data augmentation, a technique first introduced to plant disease epidemiology by [12], enables tractability of model likelihoods and so, in turn, facilitates rigorous Bayesian inference for mechanistic models. Within this framework, unobserved events, typically the timing of infection events, are treated as additional unknown parameters, and the inference is based on Markov chain Monte Carlo (MCMC) methods. These methods have had great success in analysing plant epidemic data over several years [3,9,10,13–18].

Here, we use data augmentation to fit a spatially explicit stochastic model of Flavescence dorée (FD) spread. FD is a quarantine disease threatening European vineyards. FD is one of the most damaging diseases in European vineyards. It is caused by the FD phytoplasma (taxonomic subgroups 16SrV-C and 16SrV-D), a small bacterium that multiplies in the phloem sap of infected plants. FD phytoplasma is transmitted from grapevine to grapevine by the leafhopper vector *Scaphoideus titanus* Ball [19]. Typical symptoms are leaf yellowing or reddening, downward rolling, incomplete lignification of canes, abortion of flowers, and grape wilting. FD phytoplasma, which is endemic to European alders, emerged as a disease in South West France in the 1950s, following the accidental introduction of the ampelophagous vector *S. titanus* from North America [20]. Since then, the disease has spread throughout European vineyards [21]. Due to the severe economic consequences of the disease, FD phytoplasma has been classified as a quarantine organism in Europe since 1993. There is currently no means of curing plants of FD phytoplasma. The disease is therefore controlled principally by four mandatory measures: (i) the planting of disease-free material, (ii) the application of insecticides to kill the vector, (iii) the establishment of annual vineyard surveys for monitoring plant infection, and (iv) the removal of infected plants.

The epidemiology of FD in vineyards is driven primarily by the spatio-temporal dynamics of its univoltine vector *S. titanus*. The eggs hatch in May, and there are then five nymphal instars before the first adults appear, usually in July. The adults live for about one month. The fertilised females can lay eggs from around 10 days after their emergence till the end of summer [22]. Phytoplasmas are acquired passively, through feeding on infected plants, from the first larval stage onwards. Winged adults can transmit the phytoplasma to healthy plants from early till late summer [23]. A vine plant infected during the summer of year *t* will usually become infectious from the spring of year *t* + 1. However, this infectious plant will only show the typical FD symptoms from August of year *t* + 1 [24,25]. Accordingly, FD epidemiology is characterised by the existence of cryptic infection. In the vineyard, the epidemiology of FD is also influenced by vine cultivar effects. Although no genetic resistance to FD is known, vine cultivars differ in their susceptibility to (i) phytoplasma transmission by *S. titanus* and (ii) phytoplasma multiplication and diffusion in the plant [26–28]. For example, Merlot and Cabernet Sauvignon, the two main cultivars in the Bordeaux area, display contrasting susceptibilities. Merlot is characterised by a lower proportion of infected shoots and reduced phytoplasma titers in the leaves as compared to the highly susceptible cultivar Cabernet Sauvignon [27]. These features affect FD epidemiology, as phytoplasma acquisition efficiency by *S. titanus* increases with its titers in the plant [29,30]. Direct cultivar effects on the feeding behaviour of *S. titanus* have also been shown to explain some cultivars’ higher susceptibility to FD [31].

A feature of FD epidemiology that is poorly understood is its spatial dispersal kernel. Dispersal kernels represent the statistical distribution of the location of the infected hosts after inoculum dispersal from a focal plant source [32,33]. Dispersal kernels play a crucial role in the design of control strategies for emerging pathogens that rely on reactive host removal. They consist of removing plants within a particular distance of locations identified as containing infection. The rationale is to remove (or treat) locations that are likely to be infected but not yet showing symptoms [11,33–35]. Here, we used recent georeferenced inspection and eradication data supplied by extensive surveillance at the landscape scale (representing an area of 28 hectares) to fit a mechanistic and spatio-temporal model of FD spread from one season to the next. The model builds on the work of [36] regarding the spread of Huanglongbing through citrus groves. It provides insights on the dispersal kernel that underlies FD spread. For coherent integration of information from different observations gathered at plant and field scales, we work within a Bayesian framework. We use Markov chain Monte Carlo (MCMC), a Bayesian computational method, coupled with data augmentation techniques to sample from the posterior distribution of the model parameters.

## Materials and methods

### Data

#### A plant-by-plant survey of FD symptoms in three fields

The study area is situated in Faleyras (Figure Figure S1A), a district near Bordeaux in South West France, where three plots labeled *F*_1_, *F*_2_ and *F*_3_ (Figure Figure S1B) were monitored. The initial state of the fields before the FD incursion is shown in Figure Figure S1B. In that initial state, there were 2259, 677 and 3120 plants distributed in F_1_, F_2_ and F_3_ respectively, with two different cultivars, 2354 Cabernet Sauvignon (CS) and 3702 Merlot (M), the most widely grown cultivars in the region (Fig Figure S1B). Of these cultivars, 2259 CS were planted in F_1_, 677 M in F_2_ and 3025 M and 95 CS in F_3_.

The three fields were extensively monitored on October 11, 2018, and on September 23, 2019, following the methodology described in [27]. The three fields were mapped by distinguishing the non-symptomatic, the symptomatic and the missing (due to removal) plants. Over the two studies performed by the same team (this one and [27]), in fields with high FD prevalence and sampling done at the end of summer, 98% of symptomatic Merlot plants (*n* = 207) and 99% of symptomatic Cabernet Sauvignon plants (*n* = 202) were positive for FD. Similarly, 97% of non-symptomatic Merlot plants sampled (*n* = 88) and 98% of non-symptomatic Cabernet Sauvignon plants sampled (*n* = 96) were negative in real-time PCR tests.

For each symptomatic plant, symptom severity was recorded. A severity score from 1 to 4 was assigned depending of the proportion of symptomatic branches on the stock: 1) 0 – 25%, 2) 26 – 50%, 3) 51 – 75% and 4) > 75%. The symptomatic plants were marked, and the wine grower was asked to remove them during the winter following the inspection. Note that growers may have also removed some non symptomatic plants for other agronomic reasons. The two snapshots of the health status of the fields in October 2018 and September 2019 are shown in Figure 1A and Figure 1B respectively. In 2018, 713 symptomatic plants including 668 CS and 45 M were detected, while 411 were missing (352 CS and 59 M). Field F_1_ was entirely removed during winter 2018 in compliance with FD management rules because its disease prevalence exceeded 20%. In 2019, 638 new symptomatic plants including 46 CS and 592 M were recorded on F_2_ and F_3_. During this same year, 50 additional missing plants, arising from removal of asymptomatic plants, were identified in F_2_ and F_3_.

**Fig 1.**
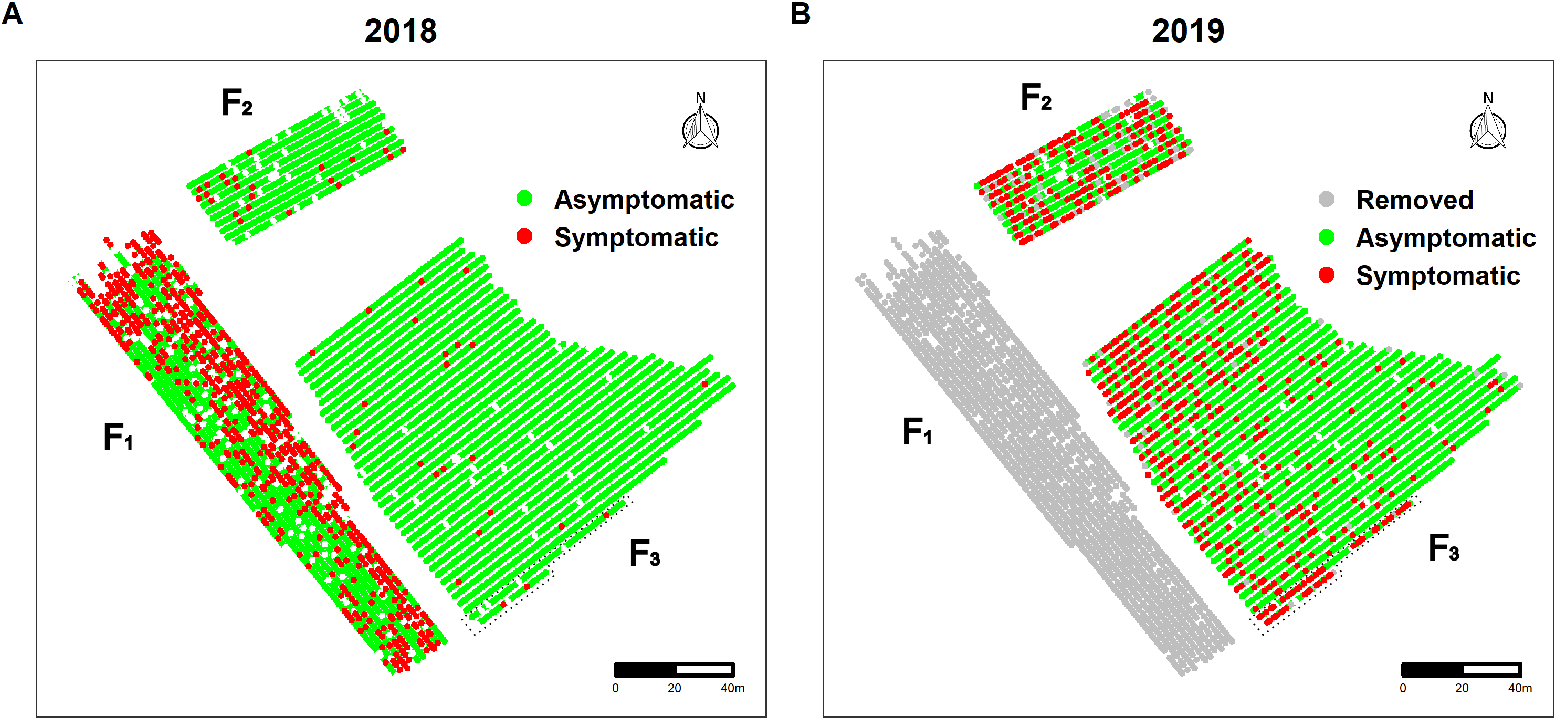
Individual status of the plants in the three vineyard fields F_1_, F_2_ and F_3_ considered. The field F_1_ was planted with 2259 Cabernet Sauvignon, the field F_2_ with 677 Merlot, and the field F_2_ with 3025 Merlot and 95 Cabernet Sauvignon in two rows surrounded by a dotted line. The status of the plants is i) asymptomatic, ii) symptomatic or iii) removed. A: Spatial distribution of 713 FD-symptomatic plants including 668 CS and 45 M detected (red) in 2018 (October 11) as well as the 411 (352 CS and 59 M) holes showing plant removals before 2018. B: Spatial distribution of 638 FD-symptomatic plants including 46 CS and 592 M detected in 2019 (September 23), 2011 plants removed in 2018 (in gray) and the same 411 (352 CS and 59 M) missing plants indicating removal before 2018. Note that real-time PCR tests indicate that more than 98% of symptomatic plants are infected with FD, while more than 97% of asymptomatic plants are FD negative.

Over two studies performed by the same team (this one and [27]), in fields with high FD prevalence and sampling done at the end of summer, 98% of symptomatic Merlot plants (*n* = 207) and 99% of symptomatic Cabernet Sauvignon plants (*n* = 202) were positive for FD. Similarly, 97% of non-symptomatic Merlot plants sampled (*n* = 88) and 98% of non-symptomatic Cabernet Sauvignon plants sampled (*n* = 96) were negative in real-time PCR tests.

#### Disease incidence in the extended area of the focal fields

The three fields, F_1_, F_2_ and F_3_, are located in a larger region where FD surveys are performed by a professional organisation known as GDON (Groupements de Défense contre les Organismes Nuisibles) des Bordeaux. The district of Faleyras was surveyed in 2014. The survey is conducted as follows. Inspectors walk around all the fields in the area in search of symptomatic plants. Once symptomatic plants are located in a given field, the inspector collects symptomatic leaves from one to five plants and pools them into a single sample. The detection of the FD phytoplasmas in samples collected is performed by accredited laboratories using a real-time PCR test [37]. Each field is then classified as infected or not. If the field is infected, the inspectors count its total number of symptomatic plants. The field will be monitored in subsequent years. The infection status of the fields monitored by the GDON des Bordeaux in 2014 within 300 m of the three focal fields is shown in Figure 2A. Note that the fields in the neighbouring district of Romagne (northeastern part of the map) were not inspected in 2014. From 2015 to 2017, the survey continued in the fields found to be infected in 2014 and spread to nearby fields after 2017 (Figure 2B-F). Note that GDON inspectors did not record the precise localisation of symptomatic plants. This information is only available in F_1_, F_2_ and F_3_ for 2018 and 2019 following a dedicated survey performed by INRAE staff.

**Fig 2.**
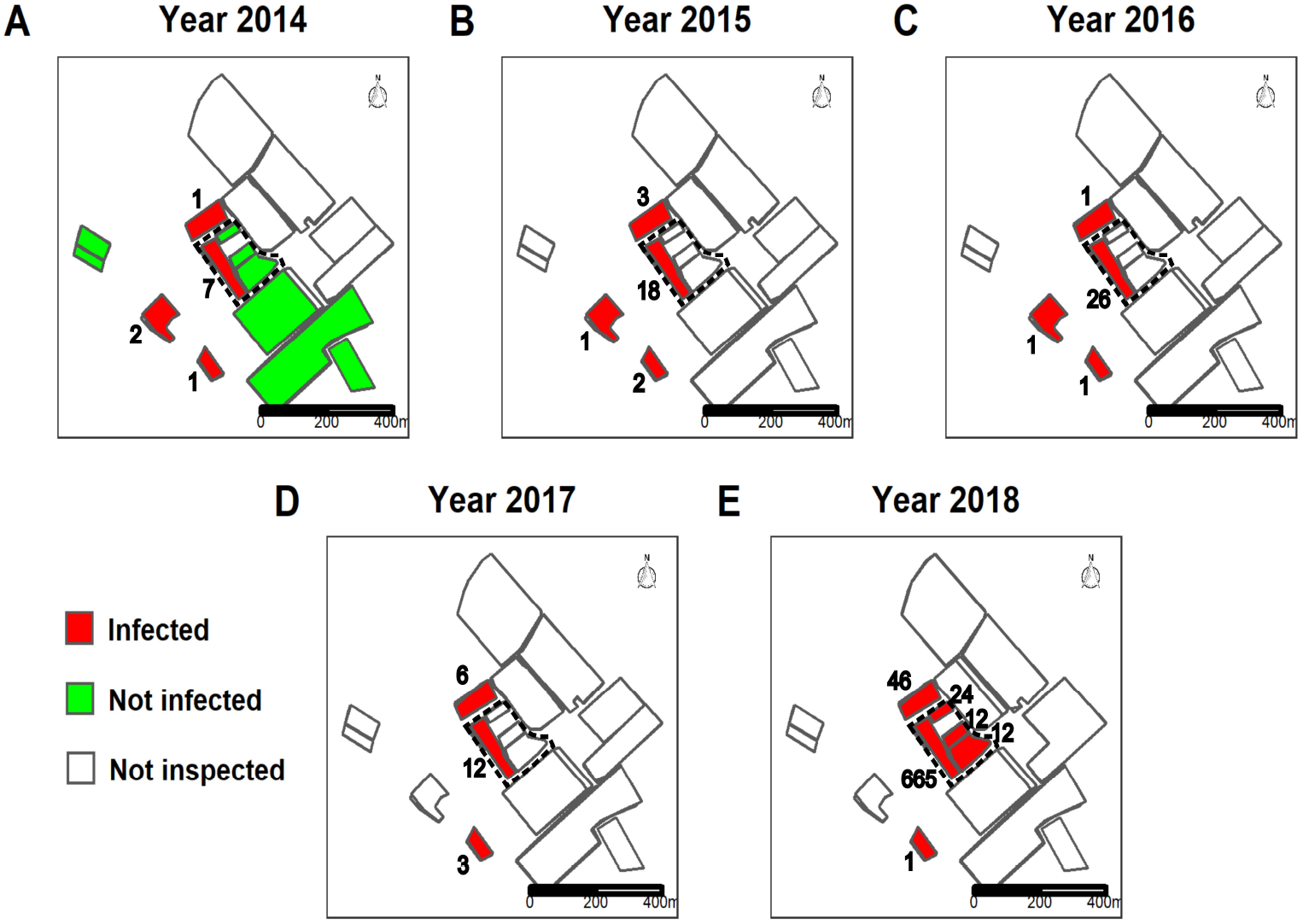
Status of fields within 300 m around the targeted fields from 2014 to 2018. The status of fields is either i) not inspected, ii) not infected by FD (if symptomatic plants were sampled but the PCR test was negative) or iii) infected by FD. For the latter category, the values inside each infected correspond to the number of symptomatic plants recorded and subsequently removed. The area with the three focal fields, F_1_, F_2_ and F_3_, is circled in dotted line.

As shown in Figure 2A, FD was first detected in the area in 2014, most prevalently in field F_1_ but also in three other fields. FD has since spread to more fields. The symptoms in the fields F_2_ and F_3_ were first reported by GDON in 2018 during the second round of inspection of F_2_ and F_3_ since 2014. Accordingly, their year of initial infection is unknown. In agreement with the set of mandatory measures to control FD, one or two insecticide treatments were recommended from 2015 to 2018 in the area corresponding to the districts of Faleyras and Romagne.

### Statistical models

We consider a mechanistic compartmental SI model with a discrete time step of one year to describe the spread of FD within the two focal fields F_2_ and F_3_. This model describes the annual infectious status of each plant from 2014 to 2019. In an SI model, plants are categorised into two infectious statuses, Susceptible (S) and Infectious (I). Susceptible plants are healthy until they become infected (i.e. infected, but not yet able to transmit infection). In FD epidemiology, the transition from susceptible to infected plants occurs during summer when adults of the vector *S. titanus* are flying. Plants infected during the summer of year *t* become infectious in year *t* + 1, meaning that they are infected and are able to transmit the infection from the start of summer of year *t* + 1. Moreover, we assume that all infectious plants are removed the following winter in compliance with the mandatory measures applied in France to manage FD.

#### Modelling the infection pressure

Let us first introduce the probability *P_i_*(*t*) that a susceptible plant *i* is infected in year *t* (and therefore becomes infectious in year *t* + 1). This probability is defined as

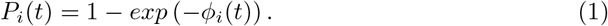

where

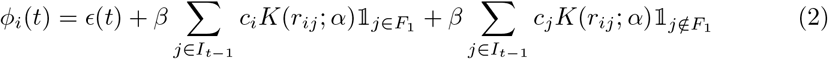

Equation 2 is the infection pressure on host *i* at year *t*. Its first two terms together represent the external infection pressure exerted on plants located in the fields F_2_ and F_3_. The first term represents the infection pressure exerted by all the fields other than F_2_ and F_3_ from 2014 to 2017, and by all the fields other than F_1_, F_2_ and F_3_ in 2018 as an infection snapshot is available for F_1_ in 2018. It corresponds to a primary infection rate that varies with time in proportion to the total number of infectious plants in the neighbouring fields (except for the plant in field F_1_ in 2018). Specifically, *ϵ*(*t*) = *ϵn*_*t*–1_, where *n*_*t*–1_ is the number of symptomatic plants at year *t* – 1 within a radius of 300 m around the two focal fields F_2_ and F_3_ (Figure 2, Table 1). The second term represents the external force of infection exerted by the infectious plants in field F_1_ in 2018 on the plants in the fields F_2_ and F_3_ in 2019. It incorporates a distance-dependent kernel *K*(*r_ij_*; *α*) shared with the third term, which represents the secondary infections taking place within the fields F_2_ and F_3_. In the second and third terms, the sum is over infectious individuals *j* (i.e. individuals infected during year *t* – 1). These terms also include a parameter *c_j_* that accounts for the differences in infectivity between cultivars. We assumed that differences in infectivity were related to the proportion of symptomatic branches on the stock. Accordingly, it was estimated independently using the severity score ranging from 1 to 4 that was described previously. Of the 668 CS plants recorded in 2018, 328 scored 1, 126 scored 2, 87 scored 3 and 127 scored 4, leading to a mean severity score value of 2.02. Of the 637 M plants recorded in 2018 and 2019, 437 scored 1, 147 scored 2, 38 scored 3 and 15 scored 4, leading to a mean severity score value of 1.42. Accordingly, we set to *c_j_* = 1.42(≈ 2.02/1.42) if the cultivar of the source plant is Cabernet Sauvignon and *c_j_* = 1 if the cultivar is Merlot.

**Table 1.** Total number of FD-symptomatic plants detected in a radius of 300 m around the focal fields F_2_ and F_3_. The values of *n_t_* include the number of symptomatic plants in all the neighbouring fields including the field F_1_ from 2014 to 2017 but excluding F_1_ in 2018. These symptomatic plants are assumed to be infectious in the model.

| t | 2014 | 2015 | 2016 | 2017 | 2018 |
| --- | --- | --- | --- | --- | --- |
| $n_t$ | 11 | 24 | 29 | 21 | 47 |

The dispersal kernel *K* is a non-negative function also known as the infection kernel. It characterises the spatial dispersal process with a parameter *α*. Specifically, it characterises the relationship between infective challenge and the relative positions of infected and susceptible plants. We propose a number of candidate models for *K*:

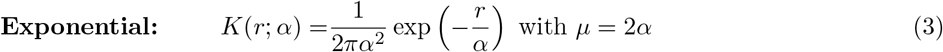

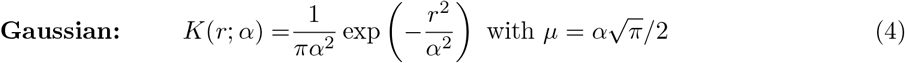

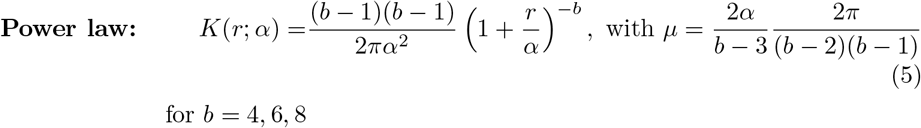

where *r* is the Euclidean distance (measured in *m*) between a given pair of plants. The mean of the dispersal kernel is given by *μ* and corresponds to the mean distance to first infection in totally susceptible population. The Gaussian and exponential kernels correspond to thin-tailed kernels, while the power law kernel corresponds to thick-tailed kernels. Kernels such as power law result in a rapid and long dispersal ahead of the source [38,39]. Note that these 1-dimensional kernels are isotropic and can represent different spatial patterns of epidemic dynamics. They also allow these different spatial patterns to be compared directly.

Overall, 20 models were fitted. They represent all combinations of two hypotheses regarding the external source *ϵ*(*i*) (1: constant, by setting to *ϵ*(*i*) = *ϵ* in equation 2, or 2: variable between years) with two hypotheses regarding a cultivar infectivity effect (1: presence, by setting *c_y_* = 1.42 for a Cabernet Sauvignon source plant and *c_y_* = 1 for a Merlot source plant, or 2: absence, by setting *c_y_* = 1 regardless of the cultivar of the source plant) and five hypotheses regarding the shape dispersal kernel. The 20 models considered are summarised in Table 2, and the corresponding equations for the infection pressure are detailed in Supplementary Information Text S1. Finally, the parameters to be estimated are summarised in Table 3.

**Table 2.** Specification of the 20 models considered. They include 2 × 2 × 5 combinations of hypotheses regarding the primary infection rate (column External source), the presence of a cultivar effect (column Cultivar effect) and the shape of the dispersal kernels (column Kernel). The numbering of equations (columns Eqn) corresponds to the equations detailed in Supplementary Information Text S1 for the infection pressure and in the main text for the kernels.

| Label | Infection pressure ($\phi(x; t)$) | | | Kernel $K(r; \alpha)$ | |
| --- | --- | --- | --- | --- | --- |
|  | External source | Cultivar effect | Eqn | Name | Eqn |
| Model 0 | Constant | No | S1 | Gaussian | 4 |
| Model 1 |  |  |  | Exponential | 3 |
| Model 2 | | | | Power law, $b = 4$ | 5 |
| Model 3 | | | | Power law, $b = 6$ | 5 |
| Model 4 | | | | Power law, $b = 8$ | 5 |
| Model 5 | Variable | No | S2 | Gaussian | 4 |
| Model 6 |  |  |  | Exponential | 3 |
| Model 7 | | | | Power law, $b = 4$ | 5 |
| Model 8 | | | | Power law, $b = 6$ | 5 |
| Model 9 | | | | Power law, $b = 8$ | 5 |
| Model 10 | Constant | Yes | S3 | Gaussian | 4 |
| Model 11 |  |  |  | Exponential | 3 |
| Model 12 | | | | Power law, $b = 4$ | 5 |
| Model 13 | | | | Power law, $b = 6$ | 5 |
| Model 14 | | | | Power law, $b = 8$ | 5 |
| Model 15 | Variable | Yes | S4 | Gaussian | 4 |
| Model 16 |  |  |  | Exponential | 3 |
| Model 17 | | | | Power law, $b = 4$ | 5 |
| Model 18 | | | | Power law, $b = 6$ | 5 |
| Model 19 | | | | Power law, $b = 8$ | 5 |

**Table 3.** Definition of model parameters and their mean posterior values along with their 95% credible interval obtained from the MCMC algorithm using Model 18. The credible interval for *t*_0_ is ignored, as there was negligible posterior weight on years other than 2016.

| Process | Symbol | Description | Estimated value | Unit<sup>1</sup> |
| --- | --- | --- | --- | --- |
| Dispersal | $\alpha$ | Scale parameter | 22.9 [20.1 - 26.5] | $m$ |
| Infection | $\epsilon$ | Primary infection rate | $10^{-4}$ [$10^{-4} - 2.10^{-4}$] | $\text{year}^{-1}$ |
| | $\beta$ | Secondary infection rate | 106.1 [94.5 - 118.4] | $m^2 \cdot \text{year}^{-1}$ |
| | $c_1, c_2$ | Cultivar’s infectivity<sup>2</sup> | $c_1 = 1$ and $c_2 = 1.42$ | na |
| Removal | $q$ | Probability of removal for reasons other than FD | 0.004 [0.004 - 0.005] | na |
| Initial infection date in $F_2 \cup F_3$ | $t_0$ | Year of first infection | 2016 | na |

<sup>1</sup> na when not applicable or for dimensionless parameters.
<sup>2</sup> Values estimated independently using data published by [27].

#### Modelling the initial infection year of the fields *F*_2_ and *F*_3_

The fields *F*_2_ and *F*_3_ were not inspected from 2015 to 2017 (Figure 2). Therefore, their unobserved levels of initial infection are treated as additional model parameters. We use *t*_0_ to denote the year the inoculum was first transmitted to either of these fields.

#### Modelling removal process

We further assume that infectious individuals are removed at the end of each year, as their symptoms are visible during summer. Finally, we assume that any susceptible individual could also be removed in a given year for some agronomic reason unrelated to FD infection. The probability *q* of this event is considered as an additional model parameter.

### Likelihood

We suppose that the data **y** record the status of plants at *n* discrete observation times in [*t*_0_, *t_max_*], along with information on removed plants. In addition, we denote the set of removed hosts with 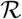, the corresponding set of reasons hosts are removed with 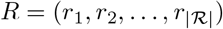 and the corresponding set of removal times with 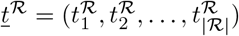. It is worth noting that 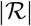 represents the size of the removed hosts in the population. For simplicity, we assume that *r_j_* is 0 if the host *j* was removed due to causes other than FD, or 1 if the host was removed due to FD infection. The set of reasons hosts are removed *R* comprises those with known reason *R_k_* and those for which the reason for removal is unknown *R_u_*. In other words, *R* = *R_k_* ∪ *R_u_*. Finally, we denote the set of the model parameters with *θ* = (*α, ϵ, β, q, t*_0_). Then, taking the reasons hosts are removed and their corresponding times as known, the complete data likelihood function is given as

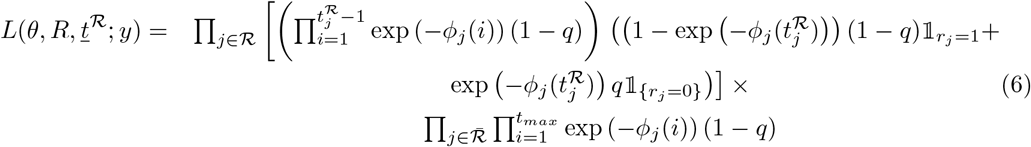

The first two lines of Eq 6 represent the contribution to the likelihood arising from the removal events. Two classes of removed hosts are considered: those removed due to the disease *r_ij_* = 1 and those removed due to causes other than FD (*r_ij_* = 0). The (1 – *q*) ensures that symptomatic plants are not removed due to any other causes. The third line represents the contribution to the likelihood of plants that escaped the removal with probability 1 – *q* and also escaped infection.

### Bayesian inference

The likelihood 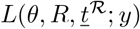 is not straightforward to compute, as the locations, dates and natures of infections between 2014 and 2017 were not observed. However, we can overcome this problem using the data-augmented approach by treating the missing plants in 2018 as an additional unknown “parameter” and investigating the joint posterior 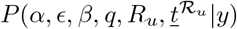 [3,13,15,17,36,40]. The algorithm involves proposing accepting changes to the times and nature of removals before 2018 and provides posterior distribution of infection sources as a by-product. This is done using methods which are now standard in computational epidemiology, such as reversible-jump [41,42] MCMC techniques. Details of algorithms are given in the electronic supplementary material (Text S1).

### Model checking and model comparison

The MCMC algorithm described in Text S1 results in a joint posterior distribution of both the model parameters and the time and nature of missing plants during the first visit to fields F_1_, F_2_ and F_3_ in 2018. These outputs can be used both to perform posterior model checking and to select the subset of candidate models that best explains the data. In the context of a spatio-temporal transmission model, natural model selection criteria such as the deviance information criterion (DIC) suffers from important limitations because of their sensitivity and complexity [43]. Following work by [12,14,15,17] in a closed context, model selection was performed by comparing (i) the posterior predictive distribution of the counts of symptomatic and removed plants and (ii) the posterior predictive distribution of the spatial structure of the epidemic to the observed ones.

#### Count of symptomatic and removed plants in 2018 and 2019

We randomly draw samples from the joint posterior distribution of the model parameters and simulate the epidemic process, assuming that the first infection(s) occurred in 2013 on the field F_1_. It is worth recalling that the initial infection on F_2_ ∪ F_3_ is estimated and therefore also sampled from the joint posterior distribution. We compared the counts of symptomatic and removed plants observed in 2018 and 2019 to those of the simulated epidemics.

#### Spatial autocorrelation

We compared the spatial structure of the simulated epidemics described above to those of the actual observations. Specifically, we considered Moran’s I index and Ripley’s L index [44–46] as our spatial summary statistics.

We divide the region into 5 × 10 square sub-regions, each containing numbers of plants ranging from 1 to 19 (with median value 16), and count the number of symptomatic sites for the time point used for the computation of Moran’s I

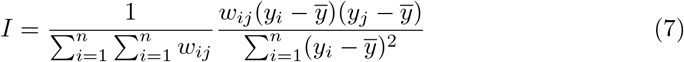

where *y_i_* = 1 if plant *i* is symptomatic before the observation time and *y_i_* = 0 otherwise, and 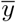 is the mean of the observation *y_i_* over *n* sub-regions. Here, *w_ij_* represents a distance-dependent weight between sub-regions *i* and *j*. There are many ways to define *w_ij_*, and here we use the distance weights between plant *i* and *j*. The index is computed by using the ape package [47] available in the statistical software R.

Similarly, we define the Ripley’s L function to characterise clustering/dispersion of point patterns at multiple distances as

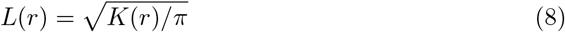

where *K*(*r*) is the Ripley K function defined as

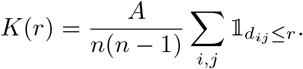

A represents the area of the window, *n* is the number of data points, and the sum is taken over all ordered pairs of points *i* and *j* in the window. *d_ij_* is the distance between the two points, and 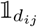 is the indicator that equals 1 if the distance is less than or equal to *r*. This spatial index is computed without requiring any aggregation of the points and therefore gives a more complete description than statistics that utilise aggregation. The index is computed by using the spatstat package available in the statistical software R [48].

## Results

### Model selection and parameter estimation

The 20 models (Table 2) were fitted to the data using reversible-jump MCMC with data augmentation (Supplementary Information Text S1, Table S1). The MCMC algorithm is iterated for 200, 000 iterations, discarding the first 50, 000 iterations (the burn-in period) to ensure the convergence to the stationary distribution is reached, with uninformative exponential priors to obtain a joint posterior distribution density for the parameters *ϵ, β, α q* and *t*_0_. Trace plots (not shown) demonstrate that there is no evidence of non-convergence across all models.

We first discuss those parameters whose posterior distributions did not differ greatly with the model fitted. First, the posterior distribution of the starting time of the epidemic reveals that 2016 was the most likely year of introduction of the inoculum into the fields *F*_2_ ∪ *F*_3_ (Figure S2). This introduction probably occurred from field F_1_, as infection in this field was first spotted in 2014. Second, we consistently estimate the 95% credible interval for the probability of plants being removed due to causes other than FD to be *q* ∈ [0.004, 0.005] per year. Third, the posterior distribution of *ϵ* (rate of primary infection from surrounding fields) depends on the assumption made (constant rate or varying with the number of symptomatic plants in the neighbourhood fields, Table 1). Nevertheless, this estimation reveals that the number of initial infections arising from primary sources is likely to be 4 for all models.

In contrast, the posterior distribution of other parameters differs significantly with the fitted models. As expected without normalizing the kernel, the infection rate β and the kernel shape *α* are highly correlated (Table S1). Normalizing the kernel by following the approach developed in [49, 50] would have reduced this correlation. The model outputs can be used to select the best model(s) by comparing the dynamics and the spatial pattern of simulated epidemics with those of the actual observation. To do so, 1000 epidemics were simulated for each model using 1000 samples from their joint posterior distributions.

The first model selection criteria compares the simulated counts of symptomatic and removed plants to observed values in 2018 and 2019 (Figures 3 and Figure S3). For clarity of these figures and similar ones, the labeling has been chosen so that each line shows the results of different parameterization of the infection pressure and each column the results of different dispersal kernels. The actual counts lie within the distribution of simulated counts only for Model 18. Models relying on Gaussian and exponential thin-tail kernels do not seem to offer an appropriate fit. Observed counts lie in the good range of the posterior count distributions provided by Models 3, 4, 8 and 9, except for the counts of removed plants in 2018.

**Fig 3.**
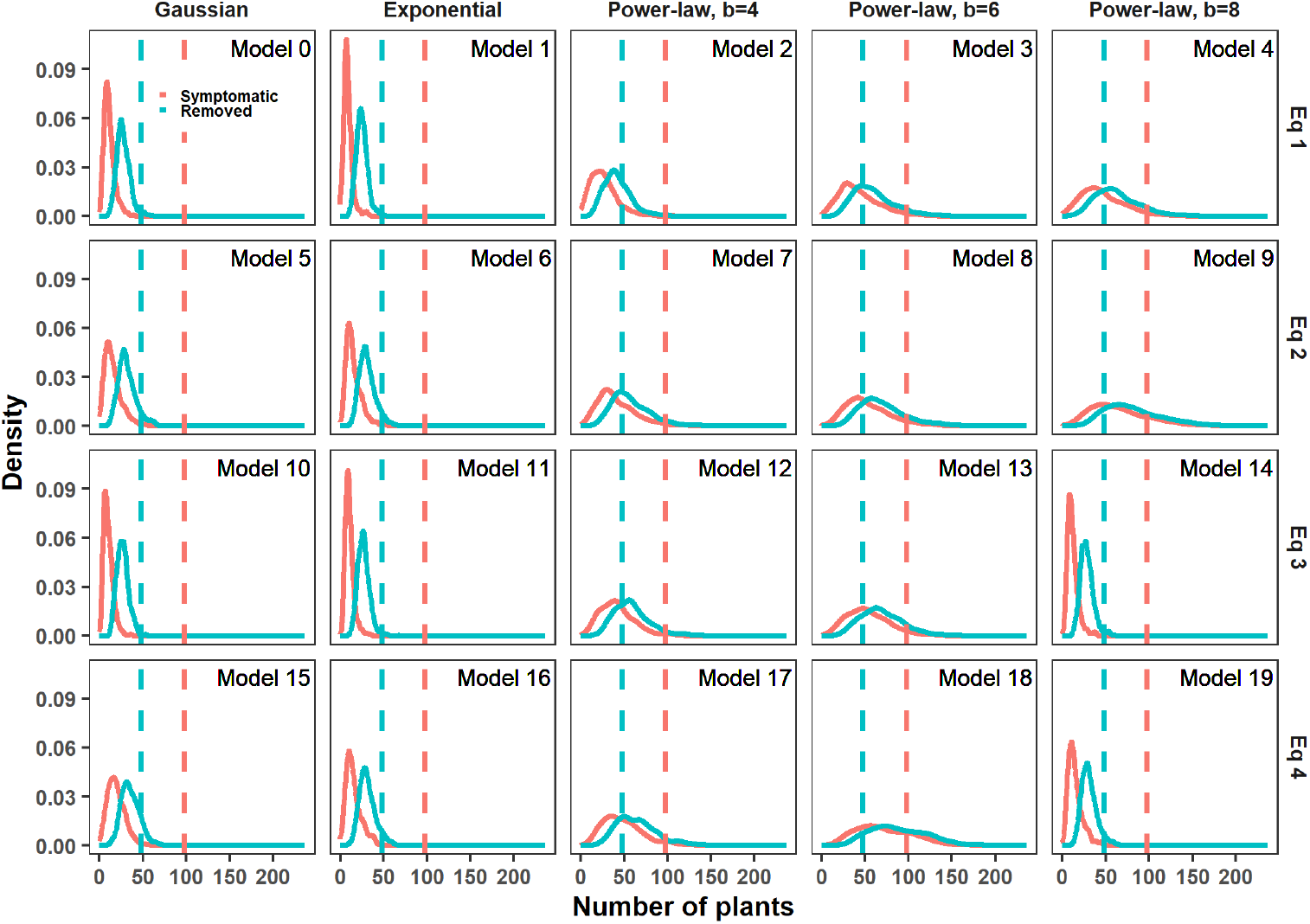
Comparison of the 20 models using the posterior predictive distribution of the counts of observed symptomatic and removed plants in 2018. Each panel corresponds to a model. The dotted line is the actual observed count, and the density of counts is obtained from 1,000 simulations. The colors correspond to the symptomatic (red) and the removed (blue) plants. The 20 models (described in Table 2) differ according to their dispersal kernel (by column) and formulation of infection pressure (by row).

**Fig 4.**
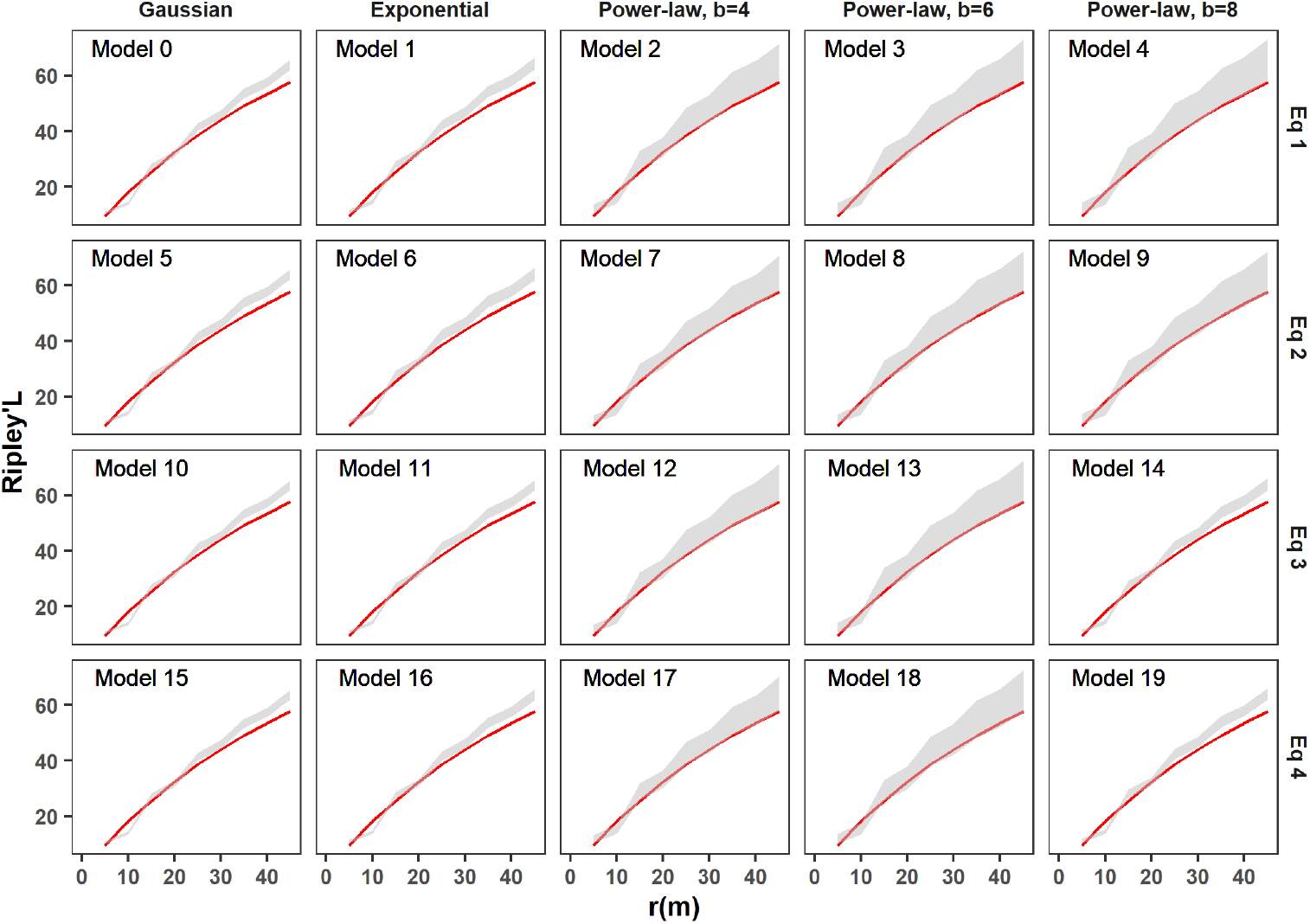
Comparison of the 20 models using the posterior predictive distribution of Ripley’s L function. The Ripley’s L function determines the clustering/dispersion of point data over a range of distances. In each panel, corresponding to a model, the red line represents the actual measure of the Ripley’s L function for the observed data, and the grey area indicates the 95% marginal credible region from the simulated epidemic. The 20 models (described in Table 2) differ in their dispersal kernel (column) and formulation of the infection pressure (row).

The second model selection criterion compares the spatial structures of the simulated epidemics to those of the actual observation using the indices of Moran’s I (Figure S4) and Ripley’s L (Figure 4). While the actual value of the Moran’s I index lies in a good range of the posterior distribution for all models, models with thin-tail kernels are completely ruled out when considering the Ripley’s L index. Power law kernel models fit reasonably well for this criterion, especially for the exponent parameters *b* = 4 and *b* = 6.

To summarise, four of the 20 models considered (Model 3, Model 4, Model 8 and Model 18) display the best fit for both observed counts and observed spatial structures. Of these, Model 18 performs slightly better. Interestingly, similar to Model 18, Models 3 and 8 rely on a power law kernel with *b* = 6. They differ by the hypothesis underlying the infection pressure, with Model 8 utilising a variable primary infection rate (but no cultivar effect) and Model 18 assuming both a variable primary infection rate and a cultivar effect, Cabernet Sauvignon being a better source of infection than the cultivar Merlot. In the following, we will focus on the results obtained from these four models, particularly Model 18.

### Dispersal kernels and FD transmission distance

A key epidemiological feature obtained in this study is the dispersal kernel of FD. It represents the rate of disease transmission between a pair of hosts at a given distance. The estimated spatial kernels are shown in Figure 5 for the 4 selected models. The posterior distribution of the kernel means *μ* shows little variation. This is evidenced by the 95% credible intervals lying within [9,18] meters (Figure 5A). In particular, the posterior mean of *μ* estimated from the best model (Model 18) is 13 meters. It is comparable to the 15 meters obtained from Model 3 and Model 8 (that only differ by the formulation of the primary infection rate). Model 4 results in a slightly lower estimate of 10 meters compared to the three others. Beside the posterior distribution of the kernel mean, we consistently observe that the rate of transmission vanishes beyond 20 meters for the 4 models considered (Figure 5B).

**Fig 5.**
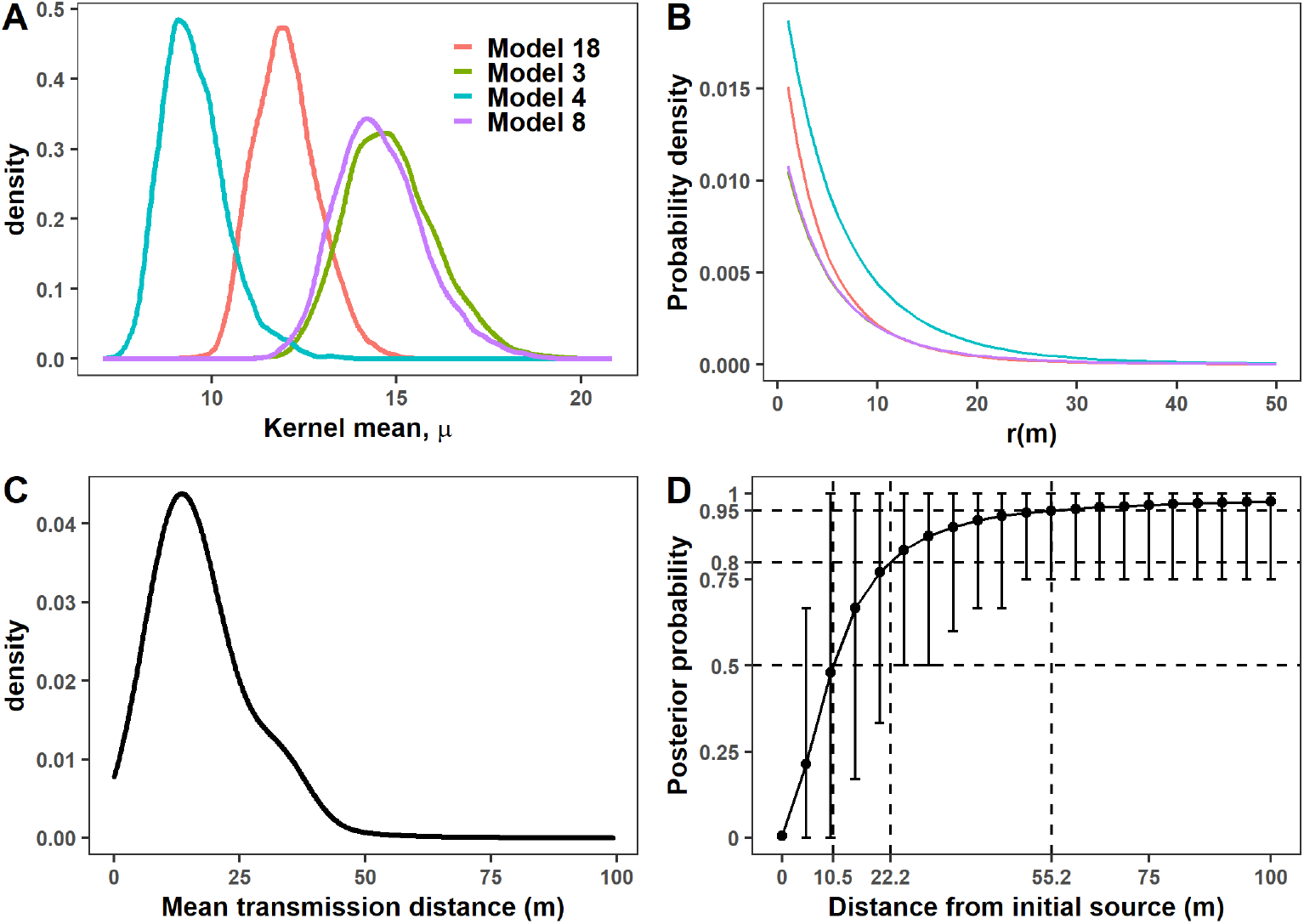
Posterior distribution of the spatial transmission under the four best models. The set of best models is Models 3, 4, 8 and 18, the last being the best one. A: Posterior distribution of the kernel mean, the mean distance to first infection in totally susceptible population. B: Posterior mean of the dispersal kernels as a function of the Euclidean distance r (in meters) between a pair of plants. C: Posterior predictive distribution of the mean transmission distance obtained by simulating epidemic realisations using 500 samples from the posterior distribution of parameters estimated from Model 18. D: Corresponding posterior predictive distribution of the cumulative distribution function (CDF) of the distance between the initial source plant and new infections in a single epidemic season (i.e. one year). Error bars represent 95% credible intervals of the CDF. The dotted lines show that on average, 50%, 80% and 95% of the new infections occur within 10.5, 22.2 and 55.2 *m* of the source, respectively.

For further insight into FD spreads, we simulate an FD epidemic in a single large square field of 1 *km*^2^ with 167334 plants. Vines are planted to mimic the practices in the Bordeaux vineyard, with spacing along rows of 3 *m* and spacing between rows of 2 *m*. For simplicity, the hosts here are assumed to be of the same cultivar so that we can ignore the effect of cultivar. We simulate disease progress over 10-year time spans, starting with a single initial infection located at the center of the field. FD spread only through secondary infections (*i.e*. no primary infection) in the considered field according to the secondary infection rate *β* and to the scale of the dispersal kernel *α* estimated from the data. We generate realisations of the epidemic process by sampling sets of parameter (*α, β*) from the joint posterior distribution of Model 18. Removal due to causes other than FD is ignored (*q* = 0), but we relax the assumption that symptomatic plants will be systematically removed one year after their infection by introducing a parameter *τ* representing the efficacy of the symptom-detection process. More precisely, the parameter *τ* corresponds to the probability that a plant infected during summer of year *t* is detected in the field survey during summer/autumn of year *t* + 1 and then removed during that winter. Viewed another way, 1 – *τ* corresponds to the proportion of infectious plants that escape removal each year, as a result of either flawed identification of symptomatic plants or flawed plant removal. Removed plants are replaced immediately. The algorithm is described in supplementary Text S1.

For each control efficiency ranging from 0 (no control) to 100% (perfect control) by steps of 20%, we run 500 realisations of the epidemic process. For each run, the observations consist of the annual snapshots of infected hosts in the 1 *km*^2^ square field considered. Specifically, we compute for each year the distances between the newly infected plants and the initial foci. Note that the secondary infection rate *β* used in the simulation is representative of the average conditions of FD spread in the area and period considered for parameter estimation. In particular, it depends on (and represents) the local population dynamics of *S. titanus* in a context where one to two annual insecticide applications are recommended (but not necessarily done, or done in suboptimal conditions).

Without a control, the localisation of infected hosts after a single cropping season allows us to estimate the posterior predictive distribution of the transmission distance, i.e. the mean distance of hosts infected from a given source in a totally susceptible population (Figure 5C). The posterior median (resp. mean) of the transmission distance is 15.01 *m* (resp. 16.9 *m*), with the 95% credible interval estimated to be [5.67, 33.59] *m*. The corresponding posterior predictive distribution of the cumulative distribution function of the distance of the newly infected plants after a single annual epidemic cycle is shown in Figure 5D. On average, 50%, 80% and 95% of the new infection occurs within 10.5, 22.2 and 55.2 *m* of the source, respectively.

The simulations also allow us to estimate how increasing control efficiency affects the distribution of the distance of the newly infected plants over 10 annual epidemic cycles. For each season and level of control efficiency τ considered, we take advantage of the 500 realisations of the epidemic to characterise the median distance, 95% quantile distance and maximum distance between the newly infected plants and the initial source plant (Figure 6 A-C). The median distance remains in the same range regardless of the level of control efficiency during the first 5 years (Figure 6A). Then, they substantially increase for low control efficiencies (*τ* ≤ 20%) while remaining in the same range for higher control efficiencies (*τ* ≥ 60%). Ten seasons after epidemic onset, infected plants are at a median distance of 273 *m* from the source without control. This distance is reduced to 212 *m* for low control efficiencies (*τ* = 20%) while remaining close to 140 *m* for control efficiencies ≥ 60%. The higher level of control efficiencies is particularly important for reducing the upper quantiles of the distance between newly infected plants of the initial source (Figure 6B,C). The maximum distance decreases almost linearly, from 604 *m* without control to 443 *m* with a control efficiency of 80%, while remaining at a close value with a perfect control. Importantly, a perfect survey/removal strategy slows down but does not stop FD spread. This is mainly due to the presence of cryptic infection. For example, ten seasons after epidemic onset, 166 *m* separate the maximum distance of localisation of newly infected plants between the no control and perfect control strategies. Beside these effects on the distributions of the distance of the new infections, increasing control efficiencies can strongly reduce the number of newly infected plants in each cropping season (Figure 6D), and thus, epidemic size. Ten years after epidemic onset, this reduction amounts to nearly 85% between the no control and the perfect strategy.

**Fig 6.**
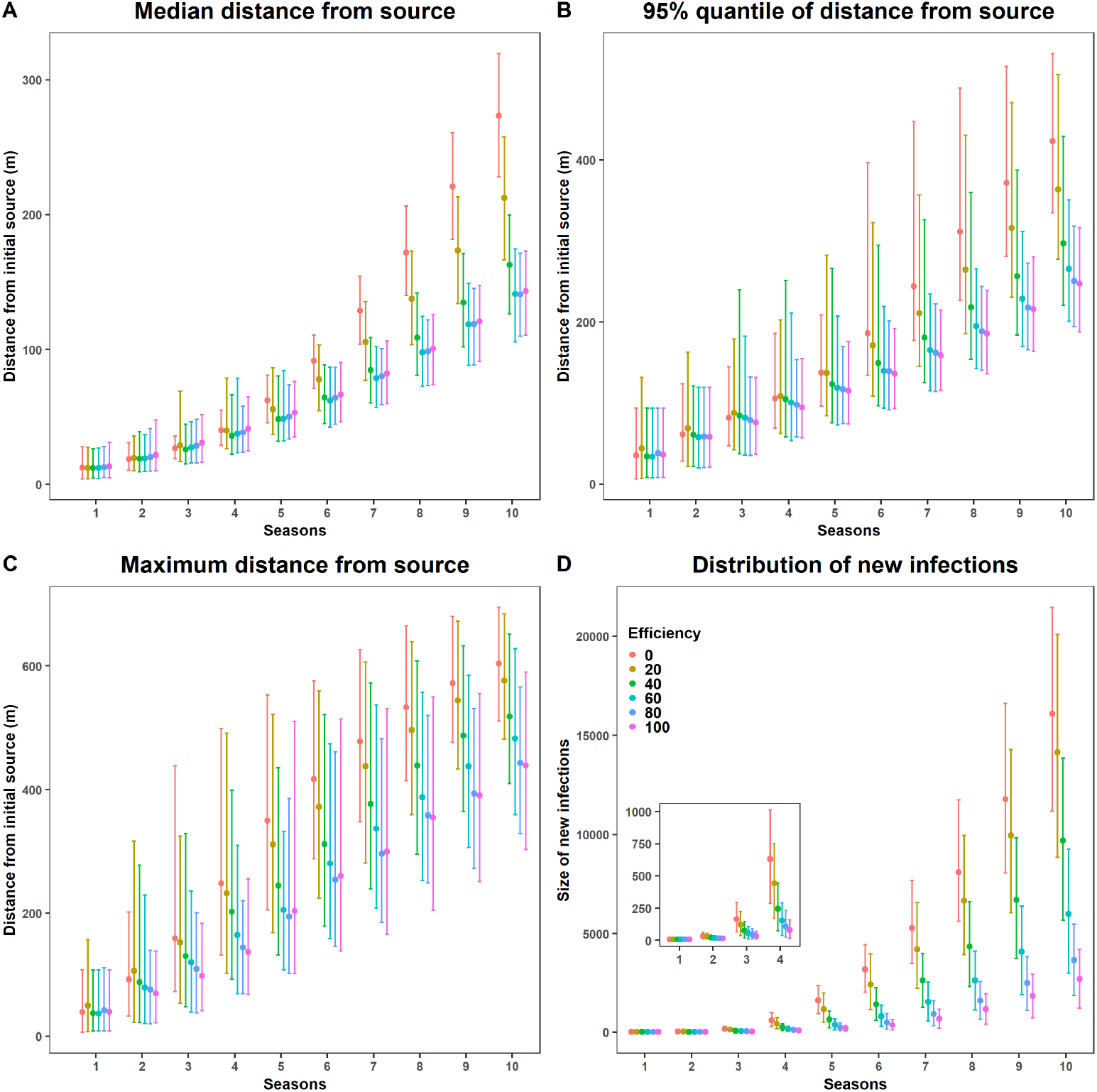
Posterior predictive distribution of distance between newly infected plants and the initial FD source and posterior predictive distribution of numbers of new infections as a function of the number of seasons after epidemic onset and control efficiency. Distance and new infections are obtained by simulating epidemics in a square field of 1 *km*^2^ with 167334 vine plants. Epidemics, initiated by a single infected plant at the center of the field, spread as a result of secondary infections while, with control, only FD-infected plants are removed and replaced. A: Posterior distribution of the median distance during 10 cropping seasons as a function of control efficiency (6 levels, from no to perfect control). B: Same as A for the 95% quantile of the distance. C: Same as A for the maximum distance. D: Posterior distributions of the mean number of newly infected plants occurring each year during 10 cropping seasons as a function of control efficiency (6 levels, from no to perfect control).

## Discussion

While estimation of dispersal parameters of plant pathogens using spatio-temporal stochastic mechanistic models has been an active research area for more than 20 years [12], we still lack information for many diseases, especially for insect-vectored pathogens [33]. In this study, we use data incorporating two consecutive annual snapshots indicating the symptomatic status of thousands of host plants into three close vineyard plots to fit a spatially explicit model representing the epidemiology of FD, a major quarantine disease damaging European vineyards. We assumed that symptom expressions reveal the infectious status of plants to FD, based on quantitative real-time PCR tests indicating that more than 98% of symptomatic (resp. more than 97% of asymptomatic) plants noted by the team in charge of the survey were positive (resp. negative) for the phytoplasma. The corresponding high sensitivity and specificity of FD detection are strongly supported for the sampling period in autumn. Indeed, the visual identification of FD symptoms is much easier then [51], as this period is the ideal time for the expression of certain specific symptoms, such as grape shrivelling and the non-lignification of canes [52]. Based on this premise, model inference relies on Markov chain Monte Carlo methods with data augmentation and allows extraction of new insights into the spread of FD. In particular, as far as we know, we provide the first characterisation of the dispersal kernel of FD. The inference was challenging in our case study because the observed data mainly consist of only two snapshots taken in two consecutive years, a situation similar to the data analysed by [12]. Such situations can be frequent for quarantine disease. Many detailed studies of plant epidemics have benefitted from more frequent spatio-temporal observations (e.g. 4 in [14], 6 in [16] and ≥ 15 in [10,17,18]). The inference framework, which reconstructs the infection dynamics in the area of interest, notably relies on the presence of missing plants (i.e. vine plants removed) before the first year of observation (2018) in the three target fields. The data augmentation methodology used here makes it feasible to infer the causes of these observed removals by considering the probability *q* that a vine plant is removed due to agronomic causes other than FD. Including this parameter allows the spread of FD during the first years after its initial incursion in the area to be slowed in a manner reflecting the situation in the field. Our results strongly suggest that the most likely date of the first infection of fields F_2_ and F_3_ is 2016 (Figure S2).

Dispersal kernels characterised by their scale parameter are the key to understanding spatial transmission of pathogens. The mean of the posterior distribution of the kernel mean *μ* obtained from the best model is 13 m (Figure 5A), and the values estimated with the 4 best models range from 10 to 15 m (Figure 5A). In addition, prediction of epidemic behaviour via simulations by sampling from the joint posterior distribution of the parameters of the best model indicates that the mean (resp. the median) of the posterior distribution of the mean transmission distance is 16.9 *m* (resp. 15.02 m) (Figure 5C). It also indicates that 50% (resp. 80%) of the new infections are likely to be found within 10.5 m (resp. 22.2 m) of the infected source plant (Figure 5D). Accordingly, FD mainly spreads locally from one year to the next. Interestingly, these estimations are in broad agreement with the space-time point pattern analysis of a FD epidemic (surveyed from 2011 to 2015) in a field located in Italy ( [53]). These authors found that the dominant feature was an aggregated cluster of symptomatic plants in moving windows of 8 × 8 plants (equivalent to a square of 11 m/side) to 12 × 12 plants (equivalent to a square of 17 m/side). They also estimated that about 50% of newly symptomatic plants (on year *n*) were found within 3 m of plants already symptomatic (on year *n* – 1) while discussing how this figure was somewhat blurred by their statistical approaches, which could not necessarily disentangle primary from secondary infections. Our estimations are also in agreement with observations on the fly activity of *S. titanus*, the vector of FD [54]. These authors have shown using mark-recapture techniques that 80% of *S. titanus* adults disperse over distances of less than 30 m along the whole season, a value in line with the aggregated spatial distribution of this monophagous leafhopper previously observed [55–57]. Some authors suggest that the crepuscular flight activity of *S. titanus* adults makes them likely to rely on an active wandering movement rather than on passive wind-borne transport [58]. More broadly, few dispersal kernels are known among insect-vectored pathogens. Huanglongbing (HLB), a major disease of citrus worldwide, is mainly transmitted by the Asian citrus psyllid, *Diaphorina citri*. The estimation of its dispersal kernel also indicates a short mean transmission distance of 5 m for 5-year-old trees and 10 m for 18-year-old trees [14]. By contrast, the mean transmission distance of plum pox virus by *Aphis gosypii* amounts to 92.8 m [10], a higher figure possibly linked to the better dispersal capacity of winged aphids that rely on wind. More broadly, it should be noted that the mean dispersal distance of fungal plant pathogens propagated by airborne spores is often much higher (ranging from 213 to 2560 m) in the four pathosystems listed by [33].

Dispersal kernels can be further defined by their shape, which in particular informs the “fatness” of their tails. Tail fatness can be used to categorise kernels in a binary fashion [38]. When at a relatively large distance the shape of the tail decreases less slowly than exponential distribution, or equally slowly, a kernel is termed “short-tailed” or “thin-tailed” [59]. In contrast, if the probability of dispersal decreases more slowly than an exponential distribution at long distances from the source, kernels are termed “long-tailed” or “fat-tailed”. Long-distance dispersal events are more frequent than with an exponential kernel that shares the same mean dispersal distance. Here, we show that the annual spread of FD is best described by a fat-tailed power law with exponent parameter *b* = 6, whereas the “thin-tailed” kernels considered (Gaussian and exponential kernels) were clearly discarded by model selection. This result is in line with the maximum distances occasionally observed for *S. titanus* by [14] (330 m) and [57] (600 m). The small-scale dispersion ability of *S. titanus* does not exclude the larger-scale passive dissemination of infectious *S. titanus* over longer distances by the wind, as is the case for many insects [60]. A consequence of fat tails is the patchiness of the epidemics that can arise. Rather than having a wave that propagates from an initial focus, you get patterns that appear to have multiple foci [12,61] and in some occasions accelerating epidemic waves [62]. These features are crucial to consider when designing control strategies of emerging plant disease [33].

Given the importance of properly characterising the speed of propagation of the front of the spread of an emerging disease, it would be interesting to consider other fat-tailed dispersal kernels, such as exponential power kernels. Indeed, fat-tailed kernels can be further distinguished depending on whether they are “regularly varying” (e.g. power law kernels) or “rapidly varying” (e.g. exponential power kernels) [59]. Mathematically, this implies that power law kernels decrease even more slowly than any exponential power function. Biologically, this means that fat-tailed exponential power kernels display rarer long-distance dispersal events than power law kernels. This affects the dynamics of an epidemic. Indeed, e.g. with weak Allee effects, subtle interactions between tail fatness (rapidly versus regularly varying) and per capita growth rate of the epidemic near zero determine whether the spread is accelerating [63,64]. However, our observations are mainly collected in three fields within a bounding box of 232 *m* * 202 *m* (i.e. over 4.7 *ha*). It is well known that data confined to relatively small spatial scales can blur the precise estimates of the form of dispersal at large distances, and in particular the shape of the kernel’s tail [65]. The selected dispersal at long distances depends on both the kernel considered and the distances over which the dataset is collected ( [66]).

The inference framework used considered a non-spatialized primary infection rate term, *ϵ*(*t*), modelling the introduction of FD in the 3 targeted fields from a set of remote fields. Recent work by [14, 17] also incorporates seasonality into the primary or external rate of infection in modelling plant epidemics, but assumes the rate to be homogeneous. Here, we utilise the data on the disease dynamic in adjacent fields by building the primary infection rate as a function of the density of symptomatic plants in those adjacent fields. This could be further refined by considering the primary infection rate as a spatially decaying source of inoculum as, in our case study, the spatial polygons of the remote fields are known (Figure 2). Thus, a further improvement of the inference framework could consist in using the same dispersal kernel for handling both the data collected at plant scale (in the fields F_1_, F_2_ and F_3_) and the data aggregated at field scale (number of symptomatic plants recorded in remote fields). Regarding this, the method developed in [67] is of interest, as it allows the quantification of the flow of particles over a heterogeneous area by integrating a pointwise dispersal function over source and target polygons. A simpler approach could also be used, e.g. through connectivity matrices between fields. However, the latter approach neglects field geometry and can be expected to bias connectivity estimates, such as when field shapes and sizes are disparate. Furthermore, given the evidence that the FD epidemic spread more rapidly along than across rows ( [53]), anisotropic kernels, in which propagules can disperse differently depending on the direction (e.g. [68]), are worth considering. A different aspect, in light of [10], that could be interesting to take into account in the inference framework is the sensitivity and specificity of FD detection (as detailed in the materials and methods section), given that we assumed here that FD detection in the focal fields was errorless.

Despite its simplicity, the spatially explicit SI model developed and fitted here captures the main features of FD epidemiology in vineyard fields: (i) a single annual infection cycle resulting from the univoltine life cycle of *S. titanus*, (ii) a latency period between summer of year *n* and spring of year *n* + 1 during which infected plants are not yet infectious and (iii) the existence of cryptic infection, since infectious plants in the spring of year *n* + 1 will only be detected beginning in August of this year (i.e. detectable symptoms strictly follow infectiousness). The simulations consider a basic scenario, all other factors impacting FD epidemiology being equal: (i) FD is spreading in a isolated and large field (100 ha) after its introduction in a single infected plant, (ii) the control consists only of removing, with the probability *τ* defining the control efficiency, infected plants as soon as symptoms become detectable (i.e. during autumn of year *n* + 1 if they were infected during year *n*) and (iii) removed plants are replanted immediately. Such control mimics the current uprooting practise in France, with the exception of the rule that states that fields must be entirely removed as soon as their FD incidences exceed 20%. This rule would be important for modelling FD control strategies in agricultural landscapes, which is out of the scope of this study. Overall, simulations suggest that even a perfect uprooting control strategy that removes all symptomatic plants (*τ* = 1) is unable to avoid FD spread (Figure 6D). It illustrates, as already emphasised in several pathosystems, that the control of emerging disease is the most difficult when it is hampered by invisible cryptic infection [11, 16]. This situation echoes the spread of FD throughout European vineyards in the last decades [21, 22] even though the FD phytoplasma has been classified as a quarantine organism since 1993. In France, the percentage of vineyards treated with insecticides against *S. titanus* increased from 50% to 75% from 2004 to 2018 (unpublished, source Grosman J. and Barthelet B., DGAL) in spite of the establishment of annual surveys for monitoring plant infection and uprooting infected plants.

Previous results should lead us to reconsider the component of the global FD control strategy that is identifying and removing infected plants. Currently, only symptomatic plants are removed in France. In the presence of cryptic infection, strategies relying on reactive host removal are of particular interest. They consist in removing plants, regardless of their infectious status (i.e. both asymptomatic and symptomatic plants), within a particular distance of infection foci. The rationale is to remove (and/or treat) locations that are likely to be infected without yet showing symptoms [11,34,35]. Note that, in line with these reasons, insecticide treatments against *S. titanus* are already advised in a buffer of 500 m around infected plots. The model developed in this work is a first step in redesigning FD control strategies. Future research should take advantage of the (S)usceptible-(E)xposed-(C)ryptic-(D)etectable-(I)nfected-(R)emoved framework proposed by [11] to optimise control strategies for invasive plant disease. Research should also benefit from one of the few attempts to model the long-term epidemiology of FD ( [69]). Using a mean-field model approximation (i.e. ignoring the transmission dispersal kernel), these authors explore the role of hotbeds (defined as vine-growing areas such as abandoned vineyards and woods containing wild grapevines) and of insecticide spray on FD dynamics. In keeping with these and our approaches, future works should consider the full spectrum of existing control measures (e.g. healthier planting material, removal of hotbeds, improving vectors with insecticides but also vibrational disturbances and “push-pull” strategies, see [22] for a review) when we only consider here the uprooting of infected plants. Finally, following [7], this approach should account for economic criteria that balance benefits (generated by the cultivation of healthy/asymptomatic plants) against costs (particularly due to FD control: surveillance, plant removal, replanting and insecticide treatments).

## Supporting information

All supplementary materials (text, table and figures)

## Supporting information

**Text S1 The supplementary text is divided into 5 sections:** (a) Equations used to model the infection pressure, (b) Description of the MCMC algorithm, (c) Parameter estimates for the 20 models fitted, (d) Model comparison and model checking, and (e) Simulation algorithm.

**Figure S1 Map of the three vineyard fields F_1_, F_2_ and F_3_ considered.**

**Figure S2 Posterior distribution of the starting year of the epidemic in fields F_2_ and F_3_ according to the 20 models fitted.**

**Figure S3 Comparison of the 20 models using the counts of observed symptomatic and removed plants in 2019.**

**Figure S4 Comparison of the 20 models using Moran’s spatial correlation index.**

**Table S1 Posterior mean, median and 95% credible region for the model parameters.**

## Data and code availability

The FD dataset, MCMC codes (written in C language) and scripts to generate the figures and tables (written in R language) are available in INRAE Dataverse at https://doi.org/10.57745/YXOEHX

## Acknowledgments

The authors wish to thank the people who sampled or provided data: Sophie Bentejac, Morgane Le Goff and Charlotte Labit (GDON des Bordeaux); Dominique Vergnes (FREDON Aquitaine); and Gontran Casella (Syndicat des Bordeaux et Bordeaux Supérieur). We also thank Sophie Bentejac and Charlotte Labit for their critical and careful reading of the manuscript. This study received financial support under research contracts Co-Act2 and RISCA (Plan National Dépérissement du Vignoble).

