## Supplementary material for "Bayesian inference for spatio-temporal stochastic transmission of plant disease in the presence of roguing: a case study to estimate the dispersal distance of Flavescence dorée": All supplementary materials (text, table and figures)

### Supplementary text S1

The supplementary text divided into 5 sections: (a) Equations used to model the infection pressure, (b) Description of the MCMC algorithm, (c) Parameter's estimates for the 20 models fitted, (d) Model comparison and model checking and (e) Simulation algorithm.

#### Equations used to model the infection pressure

Four equations were used to model the infection pressure $\phi_i(t)$ exerted on host $i$ at year $t$. They correspond to the combination of two hypotheses regarding the external source $\epsilon(t)$ (1: constant between years, or 2: variable between years) with two hypotheses regarding a cultivar infectivity effect (1: presence, by setting $c_j = 1.42$ for Cabernet-Sauvignon source plant and $c_j = 1$ for Merlot source plant or 2: absence by setting $c_j = 1$ regardless of the cultivar of the source plant).

$$\phi_i(t) = \epsilon + \beta \sum_{j\in I_{t-1}} K(r_{ij};\alpha)\mathbb{1}_{j\in F_1} + \beta \sum_{j\in I_{t-1}} K(r_{ij};\alpha)\mathbb{1}_{j\notin F_1} \quad (1)$$

$$\phi_i(t) = \epsilon(t) + \beta \sum_{j\in I_{t-1}} K(r_{ij};\alpha)\mathbb{1}_{j\in F_1} + \beta \sum_{j\in I_{t-1}} K(r_{ij};\alpha)\mathbb{1}_{j\notin F_1} \quad (2)$$

$$\phi_i(t) = \epsilon + \beta \sum_{j\in I_{t-1}} c_j K(r_{ij};\alpha)\mathbb{1}_{j\in F_1} + \beta \sum_{j\in I_{t-1}} c_j K(r_{ij};\alpha)\mathbb{1}_{j\notin F_1} \quad (3)$$

$$\phi_i(t) = \epsilon(t) + \beta \sum_{j\in I_{t-1}} c_j K(r_{ij};\alpha)\mathbb{1}_{j\in F_1} + \beta \sum_{j\in I_{t-1}} c_j K(r_{ij};\alpha)\mathbb{1}_{j\notin F_1} \quad (4)$$

#### Description of the MCMC algorithm

We employ MCMC scheme especially we use Metropolis within Gibbs sampling [1–7]. We assume vague priors for all the parameters and derive the posterior distribution of the probability of removing a plant from reasons other than FD $q$ and the joint posterior distribution of the other parameters respectively as:

$$q \sim Beta\left(1 + \sum_{j\in\mathcal{R}} \mathbb{1}_{\{r_j=0\}}, 1 + \sum_{j\in\mathcal{R}} (t_j^{\mathcal{R}} - 1) + t_{max}|\mathcal{R}| + \sum_{j\in\mathcal{R}} \mathbb{1}_{\{r_j=1\}}\right) \quad (5)$$

$$\begin{aligned} P(\alpha,\epsilon,\beta|y) = \quad & \prod_{j\in\mathcal{R}} \Big[\prod_{i=1}^{t_j^{\mathcal{R}}-1} \exp\left(-\phi_j(i)\right) \left[\left(1-\exp\left(-\phi_j(t_j^{\mathcal{R}})\right)\right)\mathbb{1}_{r_j=1} + \right. \\ & \left. \exp\left(-\phi_j(t_j^{\mathcal{R}})\right)\mathbb{1}_{\{r_j=0\}}\right]\Big] \times \\ & \prod_{j\in\bar{\mathcal{R}}} \prod_{i=1}^{t_{max}} \exp\left(-\phi_j(i)\right) \end{aligned} \quad (6)$$

**Algorithm to generate samples from the posterior distribution using Metropolis within Gibbs Sampling**

1. Initiate the chain with values $\beta^0$, $\alpha^0$, $\epsilon^0$, $q^0$, $\underline{t}^{\mathcal{R}_u^0}$ and $\mathcal{R}_u^0$.
2. Update $q$ using Gibbs Sampling by drawing from it corresponding full conditional distribution.
3. Update to $\alpha$, $\epsilon$ and $\beta$ is performed using Metropolis-Hastings

(a) Accept the new parameter $\theta_i^{new}$ with probability

$$p_{acc} = \min\left\{1, \frac{p(\theta_{\mathbf{i}}{}^{new}|\theta_{-i}, \mathbf{y})}{p(\theta_{\mathbf{i}}{}^{old}|\theta_{-i}, \mathbf{y})}\right\} \tag{7}$$

where $\theta = (\alpha, \epsilon, \beta)$ and the index $i = 1, 2, 3$ specifies the position of the corresponding parameter. $\theta_{-i}$ represents the vector of parameters exclude the parameter at position $i$.

4. Update the reason an individual was removed before 2018 using reversible-jump mcmc:

   *Repeat*

   (a) Select a host $j \in \mathcal{R}_u$, a host removed before 2018.

   (b) If $j$ was removed due to FD infection, propose to do one of the following:

   i. Move its removal date with probability 1/2. We do this by proposing a new $t_j^{\mathcal{R}^{new}}$ and accept it with probability

$$p_{acc} = \min\left\{1, \frac{L(\alpha, \beta, \epsilon, q; \underline{t}^{\mathcal{R}_u^{new}} \mathcal{R}_u^{new})}{L(\alpha, \beta, \epsilon, q, \underline{t}^{\mathcal{R}_u^{old}} \mathcal{R}_u^{old}))}\right\} \tag{8}$$

   ii. Change the reason for removal to be other reason other than infection ($r_j = 0$) with probability 1/2 i.e. the individual remains susceptible at the time of removal. We accept the new set of reasons with the probability

$$p_{acc} = \min\left\{1, 2\frac{L(\alpha, \beta, \epsilon, q; \underline{t}^{\mathcal{R}_u} \mathcal{R}_u^{new})}{L(\alpha, \beta, \epsilon, q, \underline{t}^{\mathcal{R}_u} \mathcal{R}_u^{old}))}\right\} \tag{9}$$

   (c) If $j$ was removed because of reasons other than an infection, we propose to change the reason to an FD infection. We accept the new set of reasons with the probability

$$p_{acc} = p_{acc} = \min\left\{1, \frac{1}{2}\frac{L(\alpha, \beta, \epsilon, q; \underline{t}^{\mathcal{R}_u} \mathcal{R}_u^{new})}{L(\alpha, \beta, \epsilon, q, \underline{t}^{\mathcal{R}_u} \mathcal{R}_u^{old}))}\right\} \tag{10}$$

5. Repeat i)-iv) until convergence.

#### Parameter's estimates for the 20 models fitted

For the 20 models fitted, the following parameters were estimated : (i) $\alpha$, the scale parameter of the dispersal kernel, (ii) $\epsilon$, the primary infection rate, (iii) $\beta$, the secondary infection rate, (iv) $q$, the rate of removal for reasons other than FD and (v) $t_0$, the year of first infection of fields $F_2 \cup F_3$. The estimated values of $t_0$ are given in Figure S2 and in Table S1 for all the other parameters.

**Table S1. Posterior mean, median and 95% credible region for the model parameters .**

| Parameter | 0.25% | mean | median | 97.5% | 0.25% | mean | median | 97.5% |
|---|---|---|---|---|---|---|---|---|
| | | **Model0** | | | | **Model1** | | |
| $\alpha$ | 54.440 | 60.067 | 59.897 | 66.325 | 47.180 | 55.388 | 55.146 | 65.019 |
| $\beta$ | 5.439 | 5.905 | 5.902 | 6.381 | 3.132 | 3.395 | 3.393 | 3.674 |
| $\epsilon$ | 0.001 | 0.002 | 0.002 | 0.003 | 0.001 | 0.001 | 0.001 | 0.002 |
| q | 0.004 | 0.004 | 0.004 | 0.005 | 0.004 | 0.004 | 0.004 | 0.005 |
| | | **Model2** | | | | **Model3** | | |
| $\alpha$ | 22.085 | 26.293 | 26.168 | 31.213 | 24.223 | 28.384 | 28.196 | 33.532 |
| $\beta$ | 31.078 | 34.447 | 34.388 | 38.021 | 103.737 | 116.690 | 116.549 | 130.040 |
| $\epsilon$ | 0.001 | 0.001 | 0.001 | 0.002 | 0.001 | 0.002 | 0.001 | 0.002 |
| q | 0.004 | 0.004 | 0.004 | 0.005 | 0.004 | 0.004 | 0.004 | 0.005 |
| | | **Model4** | | | | **Model5** | | |
| $\alpha$ | 25.763 | 30.305 | 29.960 | 37.149 | 53.366 | 58.839 | 58.705 | 65.239 |
| $\beta$ | 211.501 | 242.079 | 241.945 | 271.588 | 5.349 | 5.830 | 5.827 | 6.329 |
| $\epsilon$ | 0.001 | 0.002 | 0.002 | 0.003 | 0.0001 | 0.0001 | 0.0001 | 0.0002 |
| q | 0.004 | 0.004 | 0.004 | 0.005 | 0.004 | 0.004 | 0.004 | 0.005 |
| | | **Model6** | | | | **Model7** | | |
| $\alpha$ | 45.260 | 54.474 | 54.309 | 64.295 | 21.960 | 26.020 | 25.786 | 31.356 |
| $\beta$ | 3.097 | 3.363 | 3.360 | 3.649 | 30.744 | 34.161 | 34.155 | 37.622 |
| $\epsilon$ | 0.0001 | 0.0001 | 0.0001 | 0.0002 | 0.0001 | 0.0001 | 0.0001 | 0.0002 |
| q | 0.004 | 0.004 | 0.004 | 0.005 | 0.004 | 0.004 | 0.004 | 0.005 |
| | | **Model8** | | | | **Model9** | | |
| $\alpha$ | 23.880 | 27.835 | 27.626 | 33.026 | 24.760 | 28.789 | 28.558 | 34.256 |
| $\beta$ | 103.672 | 116.340 | 116.275 | 129.163 | 216.128 | 243.486 | 243.630 | 270.947 |
| $\epsilon$ | 0.0001 | 0.0001 | 0.0001 | 0.0002 | 0.0001 | 0.0001 | 0.0001 | 0.0002 |
| q | 0.004 | 0.004 | 0.004 | 0.005 | 0.004 | 0.004 | 0.004 | 0.005 |
| | | **Model10** | | | | **Model11** | | |
| $\alpha$ | 57.275 | 63.030 | 62.841 | 69.966 | 52.261 | 60.949 | 60.608 | 71.257 |
| $\beta$ | 3.902 | 4.235 | 4.232 | 4.585 | 2.266 | 2.458 | 2.457 | 2.662 |
| $\epsilon$ | 0.001 | 0.003 | 0.002 | 0.004 | 0.001 | 0.002 | 0.002 | 0.003 |
| q | 0.004 | 0.004 | 0.004 | 0.005 | 0.004 | 0.004 | 0.004 | 0.005 |
| | | **Model12** | | | | **Model13** | | |
| $\alpha$ | 18.600 | 21.939 | 21.805 | 26.112 | 20.789 | 23.872 | 23.751 | 27.748 |
| $\beta$ | 27.025 | 30.333 | 30.282 | 33.865 | 94.018 | 106.191 | 106.121 | 118.840 |
| $\epsilon$ | 0.001 | 0.002 | 0.002 | 0.003 | 0.001 | 0.002 | 0.002 | 0.003 |
| q | 0.004 | 0.004 | 0.004 | 0.005 | 0.004 | 0.004 | 0.004 | 0.005 |
| | | **Model14** | | | | **Model15** | | |
| $\alpha$ | 80.149 | 101.963 | 101.039 | 127.108 | 55.882 | 61.305 | 61.122 | 67.819 |
| $\beta$ | 103.701 | 112.734 | 112.521 | 123.013 | 3.796 | 4.152 | 4.152 | 4.510 |
| $\epsilon$ | 0.001 | 0.002 | 0.002 | 0.003 | 0.000 | 0.000 | 0.000 | 0.000 |
| q | 0.004 | 0.004 | 0.004 | 0.005 | 0.004 | 0.004 | 0.004 | 0.005 |
| | | **Model16** | | | | **Model17** | | |
| $\alpha$ | 50.114 | 59.120 | 58.757 | 69.894 | 18.437 | 21.684 | 21.567 | 25.547 |
| $\beta$ | 2.223 | 2.427 | 2.426 | 2.635 | 26.885 | 29.936 | 29.914 | 33.118 |
| $\epsilon$ | 0.0001 | 0.0001 | 0.0001 | 0.0002 | 0.0001 | 0.0001 | 0.0001 | 0.0002 |
| q | 0.004 | 0.004 | 0.004 | 0.005 | 0.004 | 0.004 | 0.004 | 0.005 |
| | | **Model18** | | | | **Model19** | | |
| $\alpha$ | 20.074 | 22.953 | 22.848 | 26.518 | 75.503 | 98.206 | 98.088 | 122.133 |
| $\beta$ | 94.462 | 106.099 | 105.959 | 118.383 | 101.944 | 111.427 | 111.334 | 121.829 |
| $\epsilon$ | 0.0001 | 0.0001 | 0.0001 | 0.0002 | 0.0001 | 0.0001 | 0.0001 | 0.0002 |
| q | 0.004 | 0.004 | 0.004 | 0.005 | 0.004 | 0.004 | 0.004 | 0.005 |

#### Model comparison and model checking

The model selection was performed by comparing simulated (i) counts of symptomatic and removed plants to the observed ones and (ii) the spatial structure of the epidemic. The supplementary text displays the comparison of simulated and observed counts of symptomatic and removed plants in 2019 (Figure S3) and the comparison of the spatial structure of the observed and simulation epidemics based on the Moran's I index (Figure S4).

#### Simulation algorithm

**FD simulation algorithm**

1. Initialise $t = t_0$, the initial time, and set the location of the initial infection to be $X_0 = (x_0, y_0)$, the number of infectious hosts $n = 1$.
2. New infections (or removals) are generated as follows. Suppose that at time $t$ there have been $k$ active infections and let $P_j(t+1)$ the probability that a susceptible plant $j$ gets infected at year $t+1$. Recall from the main text that

$$P_j(t) = 1 - exp\left(-\phi_j(t)\right). \tag{11}$$

where

$$\phi_j(t) = \beta \sum_{i \in I_{t-1}} K(x, y; \alpha) \tag{12}$$

Draw $u_1 \sim U(0,1)$ and $u_2 \sim U(0,1)$

3. If $u_1 < P_j(t+1)$, then the plant at position $x$ is infected else it remains susceptible and set $t = t + 1$.
4. If $u_2 < \tau$ for any of the $k$ active infected plants, remove such a plant and replace it with a new healthy plant and set $t = t + 1$. Recall that $\tau$ represents the efficiency of the removal.
5. Repeat $2 - 4$ until a stopping criterion is reached (e.g. $t \geq T$).

#### Supplementary figure S1

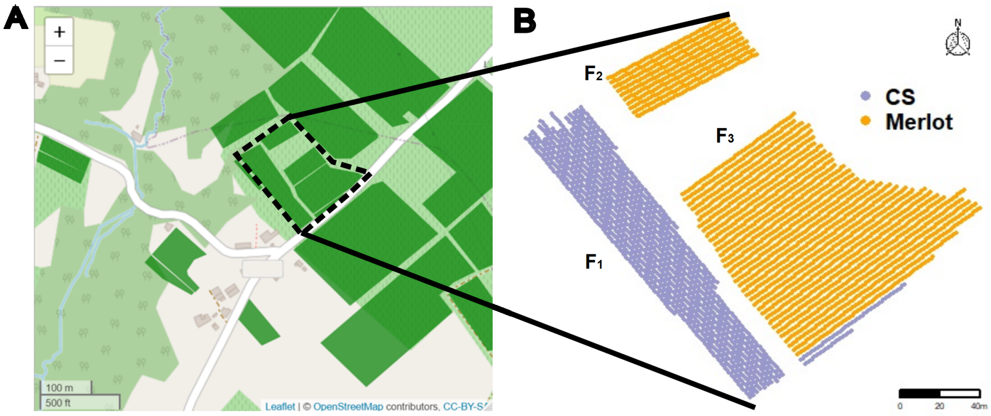


**Fig S1. Map of the three vineyard fields $F_1$, $F_2$ and $F_3$ considered.** A: Map of the study area in Faleyras, a district in South West France. The map shows the targeted fields (highlighted in the box) and the neighbouring fields within 300 m. B: Initial state of the fields where 2259 Cabernet Sauvignon (CS) were planted in field $F_1$, 677 Merlot in field $F_2$, and 3025 Merlot and 95 Cabernet Sauvignon in field $F_3$.

### Supplementary figure S2

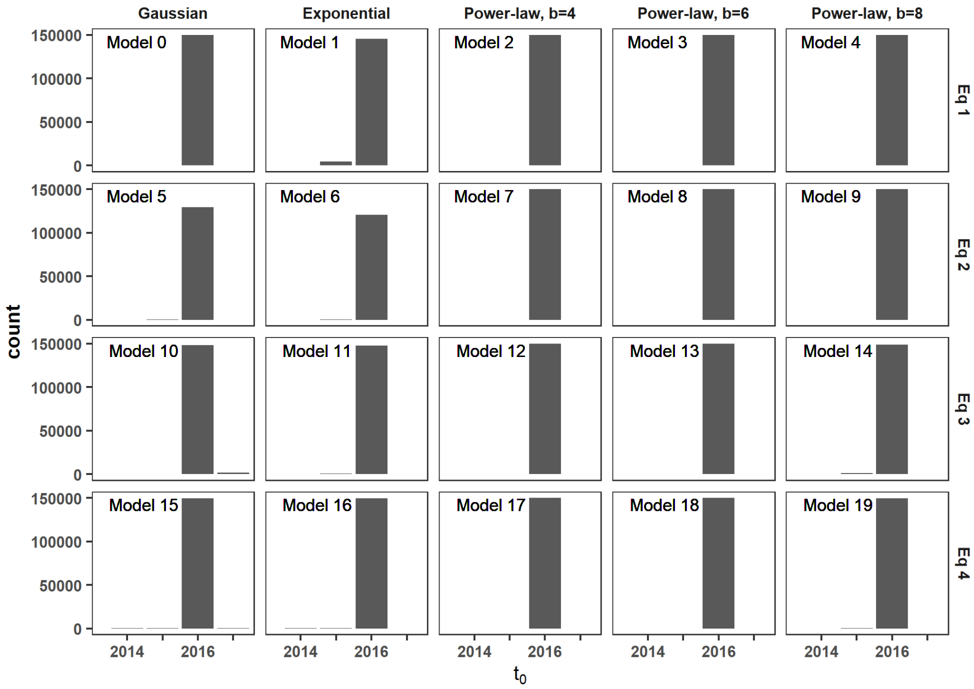


**Fig S2. Posterior distribution of the starting year of the epidemic in fields $F_2$ and $F_3$ according to the 20 models fitted.** The 20 models differ according to their dispersal kernel (in column) and formulation of the infection pressure (in row).

#### Supplementary figure S3

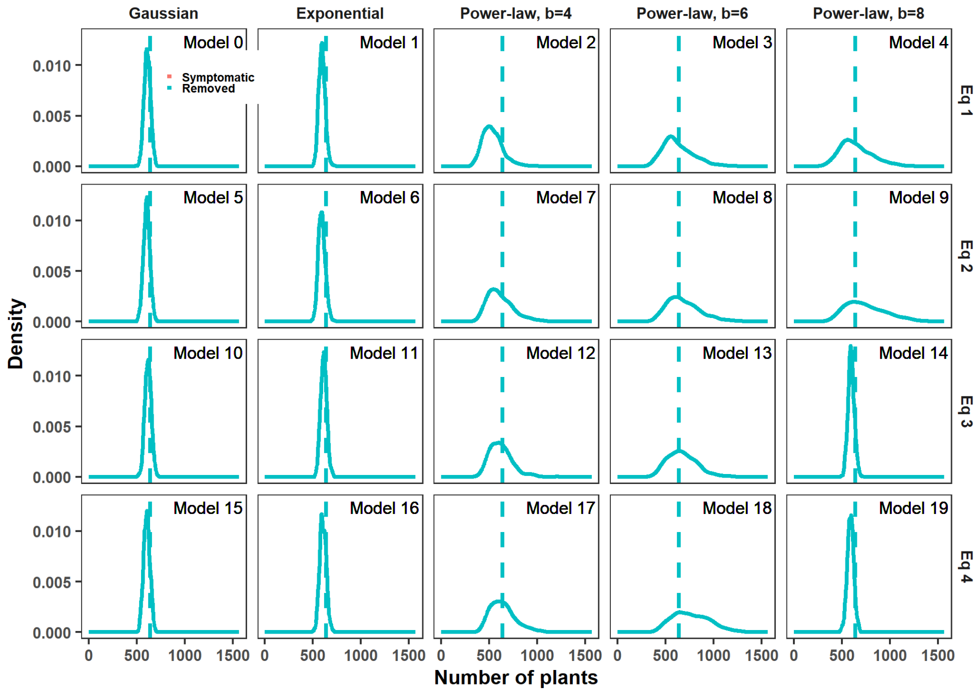


**Fig S3. Comparison of the 20 models using the counts of observed symptomatic and removed plants in 2019.** In each panel, corresponding to a model, the dot line is the actual observed count and the density of counts are obtained from 1000 simulations. The colors correspond to the symptomatic (red) and the removed (blue) plants. The 20 models differ according to their dispersal kernel (in column) and formulation of the infection pressure (in row). Symptomatic and removed plants in 2019 coincident here since we did not allow for removals for reasons other than FD in 2019 as this information was not available. In fact, this should have been available on the 2020 snapshot.

### Supplementary figure S4

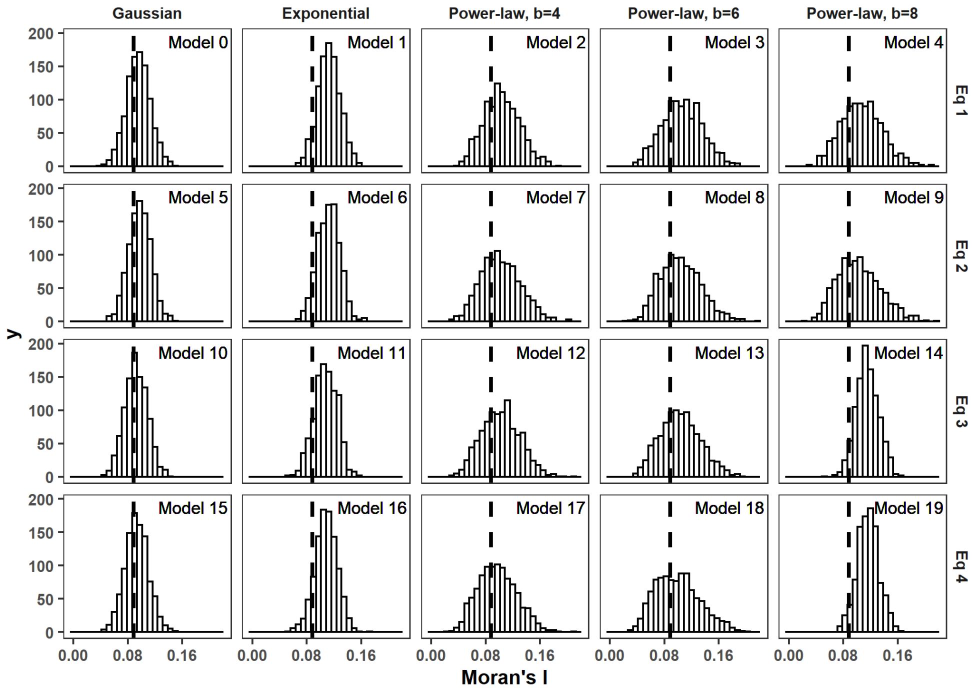


**Fig S4. Comparison of the 20 models using the spatial correlation index of Moran.** Posterior predictive distributions of spatial correlation using Moran's I index [8]. In each panel, corresponding to a model, the black line represents the actual measure for the observed data. The 20 models differ according to their dispersal kernel (in column) and formulation of the infection pressure (in row).
